# Adaptation of proteins to the cold in Antarctic fish: A role for Methionine?

**DOI:** 10.1101/388900

**Authors:** Camille Berthelot, Jane Clarke, Thomas Desvignes, H. William Detrich, Paul Flicek, Lloyd S. Peck, Michael Peters, John H. Postlethwait, Melody S. Clark

**Author notes:** Corresponding Author: Melody S Clark, British Antarctic Survey, Natural Environment Research Council, High Cross, Madingley Road, Cambridge, CB3 0ET, UK.

## Abstract

The evolution of antifreeze glycoproteins has enabled notothenioid fish to flourish in the freezing waters of the Southern Ocean. Whilst successful at the biodiversity level to life in the cold, paradoxically at the cellular level these stenothermal animals have problems producing, folding and degrading proteins at their ambient temperatures of down to −1.86°C. In this first multi-species transcriptome comparison of the amino acid composition of notothenioid proteins with temperate teleost proteins, we show that, unlike psychrophilic bacteria, Antarctic fish provide little evidence for the mass alteration of protein amino acid composition to enhance protein folding and reduce protein denaturation in the cold. The exception was the significant over-representation of positions where leucine in temperate fish proteins was replaced by methionine in the notothenioid orthologues. Although methionine may increase stability in critical proteins, we hypothesise that a more likely explanation for the extra methionines is that they have been preferentially assimilated into the genome because they act as redox sensors. This redox hypothesis is supported by the enrichment of duplicated genes within the notothenioid transcriptomes which centre around Mapk signalling, a major pathway in the cellular cascades associated with responses to environmental stress. Whilst notothenioid fish show cold-associated problems with protein homeostasis, they may have modified only a selected number of biochemical pathways to work efficiently below 0°C. Even a slight warming of the Southern Ocean might disrupt the critical functions of this handful of key pathways with considerable impacts for the functioning of this ecosystem in the future.

## Introduction

One of the many consequences of our warming world will be the irretrievable loss of the coldest habitats. This change will have a significant impact on the endemic fauna, many of which have evolved novel adaptations to life in freezing conditions (Pörtner et al. 2007). Clearly, there is strong interest in learning how these cold-adapted species will respond in the coming years, in particular with regard to those species of economic importance such as the notothenioid fish in the Southern Ocean. We are constrained in our abilities to decipher responses to change, however, because we know relatively little about the molecular genetic mechanisms these species have evolved to thrive in freezing conditions and how these adaptations will impact their future responses and resilience in a changing world.

Rapid climate change is affecting oceans in both polar regions, more specifically the Arctic and along the Antarctic Peninsula (Nicholas et al. 2014; Abram et al. 2014). Although both regions contain freezing seas, their faunas have different evolutionary histories based on geography (Hunt et al. 2016). The North Pole is in the open ocean and Arctic benthic fauna are largely a product of the Pleistocene glaciation (2.58 MYr) and with few endemic species (Dunton 1992). In comparison, the break-up of Gondwana resulted in the South Pole being in the middle of a continent. The opening of the Drake Passage (25-22 MYr) between South America and Antarctica, and the subsequent development of the Antarctic Circumpolar Current around the continent and the Antarctic Polar Front (Maldonado et al. 2014; Sher et al. 2015) effectively isolated the marine fauna around Antarctica. The Southern Ocean gradually cooled with the formation of sea ice 14-12 MYr ago (Shevenell et al. 2011). The resident fauna has, therefore, been subjected to long and intense selective pressure for survival in freezing waters. In general, Antarctic species are characterised by long development and generation times, deferred maturity and extended life spans (Peck 2018); thus, a key factor to which they are not adapted is rapid environmental change.

These evolutionary pressures of isolation and cooling in the Southern Ocean led to local extinction of most fish fauna but a massive radiation of the notothenioid fish, which are classed as a rare example of a marine species flock (Lecointre et al. 2013). Of 129 catalogued notothenioid species, 101 are present only in the Southern Ocean (Eastman and Eakin 2000) and key members of this order are targets for commercial fisheries. Notothenioids have developed some very distinct adaptations to life in the cold and the associated characteristic of highly oxygenated waters (Carpenter 1966; Peck 2018). Generalised molecular adaptations include the possession of antifreeze glycoproteins (Devries 1971; Chen et al. 1997; Near et al. 2012), cold-stable yet dynamic cytoplasmic microtubules (Billger et al. 1994; Detrich et al. 1989, 2000; Himes and Detrich 1989), the protein-folding complex CCT (Cuellar et al. 2014), giant muscle fibres with reduced fibre numbers (Johnson et al. 2003), and the loss of a typical inducible heat shock response (Hofmann et al. 2000; Clark et al. 2008). One family, the Channichthyidae (commonly known as the icefish), are highly derived – they lack haemoglobin and functional erythrocytes and several icefish species are devoid of myoglobin in both cardiac and skeletal muscle (Moylan and Sidell 2000; di Prisco et al. 2002).

Whereas protein denaturation at high temperatures has long been recognized, there is less appreciation that proteins denature in response to cold stress (Privalov 1990; Peck 2016). At the molecular level, a number of studies have identified amino acid changes that increase the flexibility of some proteins, allowing them to work more efficiently around 0°C (Capasso et al. 1999; Detrich et al. 2000; Roach 2002; Xu et al. 2008; Cuellar et al. 2014). Other results have shown that retention of gene duplicates by cold-living species helps to ensure that protein production levels are maintained at levels comparable to temperate species, albeit using twice the number of genes (Carginale et al. 2004; de Luca et al. 2009). These studies, however, are limited in number and may not represent global protein metabolism in Antarctic species. Other reports indicate that protein production, un-folding and accumulation of ubiquitinated proteins may be significant problems in Antarctic species (Peck 2016, 2018). Antarctic fish contain higher levels of ubiquitin-tagged proteins than closely related temperate species and constitutively express the “inducible” form of the 70-kDa heat shock protein (HSP70), a molecular chaperone that helps to rescue denatured proteins (Hofmann et al. 2000; Place et al. 2004; Place and Hofmann 2005; Clark et al. 2008; Todgham et al. 2007, 2017). Thus, steady-state protein production and functioning is likely to be much less efficient in Antarctic species compared with temperate relatives, despite the protein adaptations detailed above. Identification of the extent to which the proteome of Antarctic notothenioid fish is cold-adapted should provide critical information for predicting how these species will respond to a changing world. Fortunately, our ability to tackle these questions by interrogating protein amino acid sequences for cold-adapted substitutions is improving rapidly.

Since the original Antarctic toothfish *Dissostichus mawsoni* study on Expressed Sequence Tags (ESTs) (Chen et al. 2008) and the emergence of Next Generation Sequencing, molecular data for notothenioids has gradually increased. These data have largely been obtained using Roche 454 with published studies often restricted to a single species (Windisch et al. 2012; Shin et al. 2012; Coppe et al. 2012; Papetti et al. 2015). To date, only head kidney tissue has been subjected to short read Illumina sequencing (Gerdol et al. 2015), with a mixed approach used to generate the first draft genome of the bullhead notothen *Notothenia coriiceps* (Shin et al. 2014). Three studies have developed preliminary transcriptomes with the aim of identifying notothenioid responses to warming (Huth and Place 2013; Bilyk and Cheng 2013, 2014), but given the piecemeal information available with regard to tissues, treatments and different sequencing platforms, it is difficult to directly compare results across studies to generate an unbiased global overview of notothenioid gene evolution to the cold.

In this study, we sequenced the transcriptomes of four Antarctic notothenioids using high-throughput sequencing. These species included two icefish (*Neopagetopsis ionah* (Jonah’s icefish) and *Pseudochaenichtys georgianus* (South Georgia icefish)) and two red-blooded species (*Harpagifer antarcticus* (Antarctic spiny plunderfish) and *Parachaenichthys charcoti* (Charcot’s dragonfish)) (Figure 1). These species are phylogenetically distinct within the notothenioids, being taxonomically assigned to three families (Channichthyidae, Harpagiferidae and Bathydraconidae) and four genera (Near et al. 2012). These species choices enabled us to differentiate between generalised notothenioid-specific changes in amino acid composition and those that are species-specific in comparison to orthologous sequences from temperate teleost relatives. These global analyses are described and discussed in the context of cold-adaptation and the hypothesis that Antarctic fish proteomes are incompletely adapted to function efficiently in freezing oceans.

**Figure 1.**
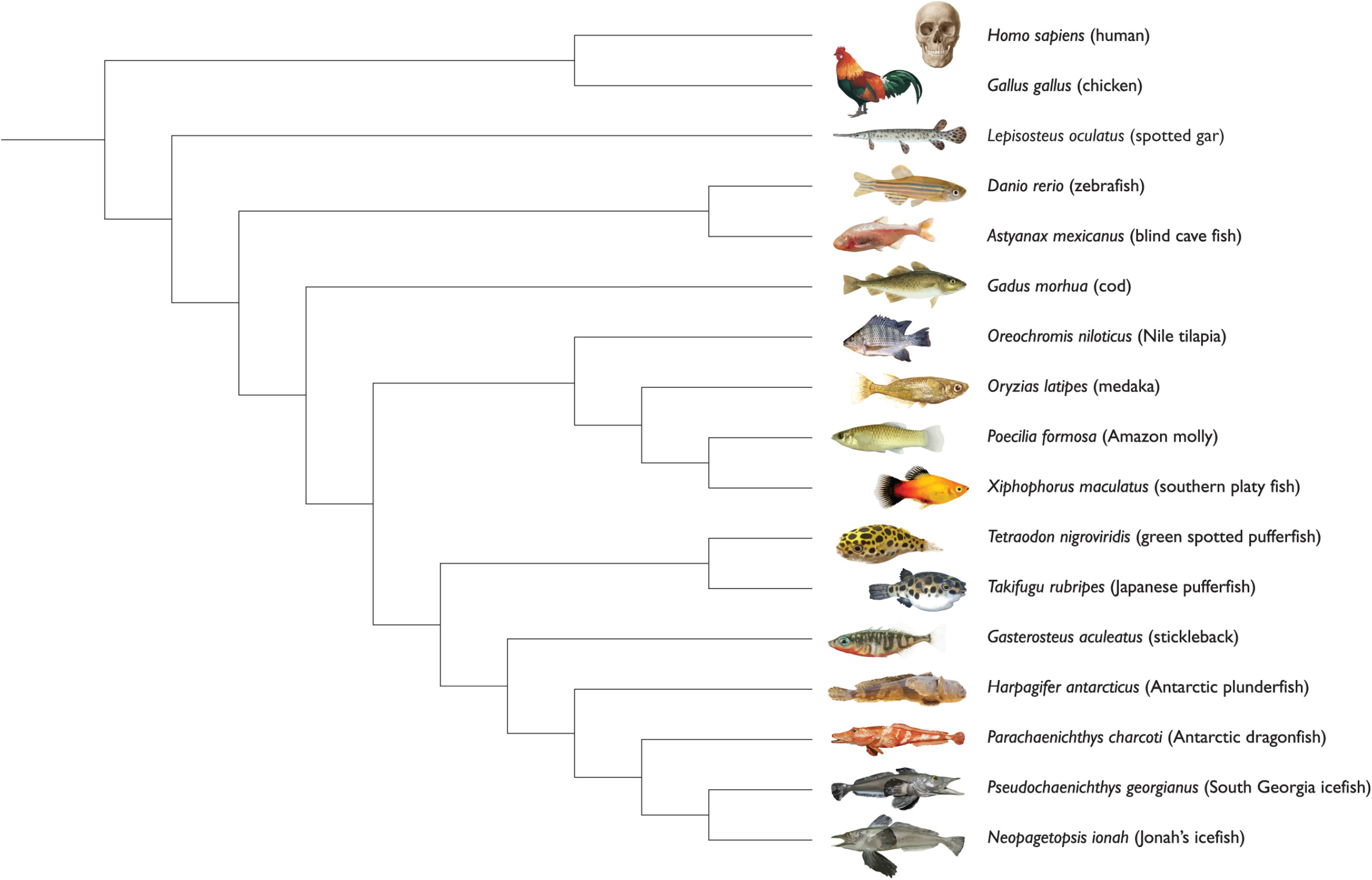
Phylogenetic placement of notothenioid fish to demonstrate relatedness to other species used in these analyses and for the positive selection scans. Based on Near et al. (2012) and Betancur et al. (2013).

## Results

### Transcriptome assemblies

To document the protein-coding gene content of notothenioid genomes, we performed mRNA-seq on one to four individuals for each of the four species included in the study. Samples were collected from animals captured in Ryder Bay, south of Low Island, west of Brabant Island and in Andvord Bay during field trips in Antarctica in 2013 and 2014 (Figure 2). Tissue collection was driven by availability and included one to thirteen tissues per individual (Supplementary Table S1). As a result, the obtained transcriptomes varied in quality and completeness across species: *P. georgianus* and *P. charcoti* were the most exhaustively sampled, while *N. ionah* was only represented by one tissue (spleen) from one individual. Sequencing yielded from 8 to 39 million paired-end reads per library, which constituted a total of 14 to 503 million read pairs per species.

**Figure 2.**
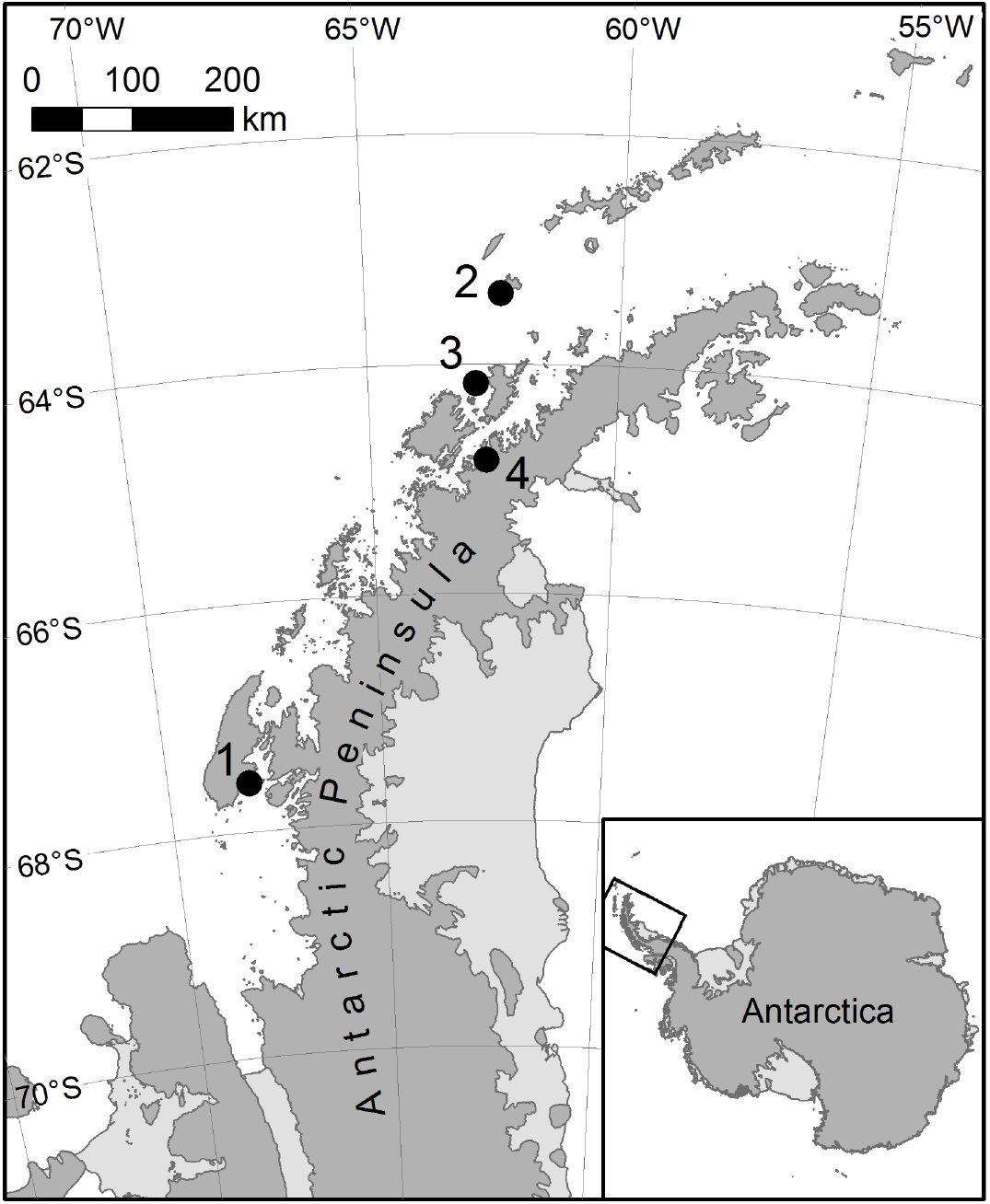
Wild capture locations for the notothenioid fish samples used in this study. Numbers correspond to the following locations: 1: Ryder Bay, 2: south of Low Island (Antarctic Specially Protected Area No. 152, Western Bransfield Strait), 3: west of Brabant Island (Antarctic Specially Protected Area 153, Eastern Dallmann Bay) in the Palmer Archipelago, 4: Andvord Bay near Palmer Station

Transcriptomes were assembled *de novo* using Trinity (Grabherr et al. 2011) on quality-filtered pooled reads from all individuals and tissues for each notothenioid species (Methods). The assemblies contained 72,573 to 251,528 transcripts across species, grouped into 61,696 to 212,979 ‘gene’ units by Trinity (Table 1). Median contig length was ~400 bp and was consistent across species. Over 80% of initial reads could be realigned to the assembled transcriptomes, which is consistent with high assembly quality (Hass et al. 2013). We then assessed transcriptome completeness using the Benchmarking Universal Single-Copy Orthologs (BUSCO) method (Simão et al. 2015). This analysis confirmed that the great majority of single-copy genes found across all vertebrate and Actinopterygian species were present and complete in our assemblies (Figure 3). As expected, the *N. ionah* transcriptome from a single tissue was less exhaustive than for the other three species; nonetheless, 62% of reference Actinopterygian genes were successfully identified as full-length transcripts in this species.

**Table 1.** *De novo* assembly and annotation statistics for the four notothenioid transcriptomes included in this study. ‘Gene’ units correspond to Trinity’s definition of a gene, i.e. a group of transcripts that likely are isoforms from the same gene.

|  | *P. georgianus* | *P. charcoti* | *H. antarcticus* | *N. ionah* |
| --- | --- | --- | --- | --- |
| Number of sampled individuals | 4 | 2 | 1 | 1 |
| Number of sampled tissues | 10 | 13 | 6 | 1 |
| Total number of mRNA-seq libraries | 28 | 23 | 6 | 1 |
| Total pairs of paired-end reads | 503,200,884 | 368,989,424 | 89,734,790 | 14,297,063 |
| Number of transcripts in *de novo* assembly | 251,528 | 173,461 | 145,576 | 72,573 |
| Number of 'gene' units | 212,979 | 144,437 | 118,095 | 61,696 |
| Median transcript length (bp) | 374 | 390 | 408 | 442 |
| Mean transcript length (bp) | 822 | 914 | 890 | 874 |
| N50 (bp) | 1,725 | 2,032 | 1,827 | 1,624 |
| Total assembled bases | 206,860,807 | 158,515,494 | 129,565,375 | 63,417,105 |
| Fraction of reads mapping to transcripts | 83.8% | 80.0% | 80.7% | 81.7% |
| Number of transcripts with temperate orthologs | 26,072 | 26,974 | 24,963 | 16,941 |
| Number of genes in orthology groups | 10,728 | 10,728 | 8,611 | 7,093 |

**Figure 3.**
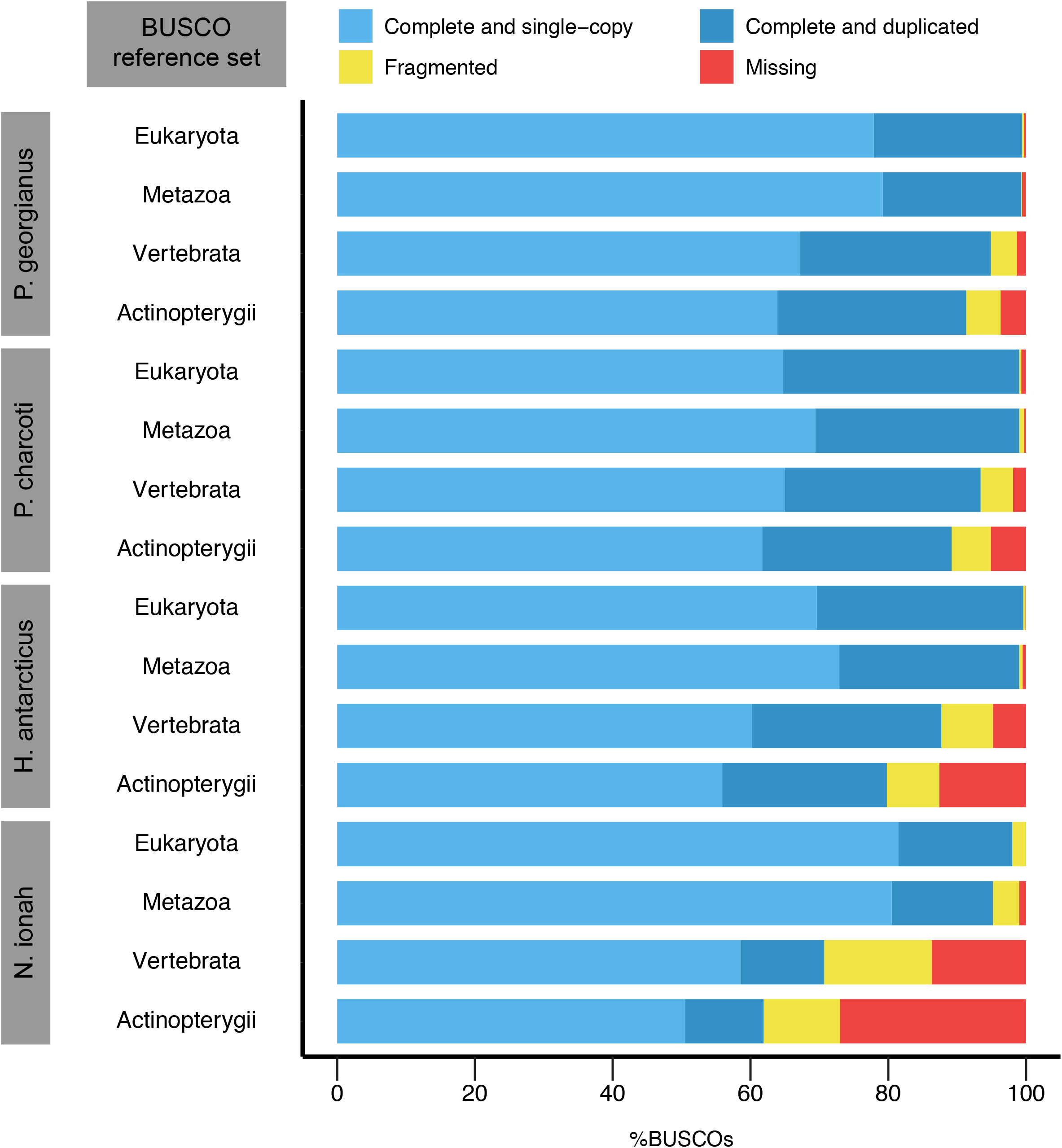
Completeness assessment of the four de novo notothenioid transcriptome assemblies using the BUSCO (Simão et al. 2015) reference gene sets. The reference sets include 303, 978, 2,586 and 4,584 genes for *Eukaryota, Metazoa, Vertebrata* and *Actinopterygii* respectively.

### Identification of orthologous genes

We used stickleback (*Gasterosteus aculeatus*) and zebrafish (*Danio rerio*) as high-quality temperate teleost reference genomes to identify and annotate orthologous sequences across notothenioid transcriptomes. Stickleback is the species with a sequenced genome that is both most closely related to notothenioids (Figure 1) and for which there is good annotation and functional validation of gene roles. Because our proposed analyses relied on accurate rather than exhaustive orthology identification, we used a fairly conservative approach. Briefly, notothenioid transcripts were compared to stickleback and zebrafish transcriptomes using BLAST (Altschul et al. 1990) with stringent e-value thresholds to identify the top match as a putative orthologue in each reference species (Methods). We considered the orthology relationship as high-confidence when the putative orthologues in stickleback and zebrafish correspond to an orthologous gene pair in the Ensembl database (Vilella et al. 2009; Herrero et al. 2016). We successfully identified 16,941 to 26,974 transcripts with high-confidence orthologues in temperate fish across notothenioid transcriptomes (Table 1). This process resulted in the identification of 10,728 fully consistent orthology groups present in stickleback, zebrafish and at least two of the notothenioid fish (*P. georgianus* and *P. charcoti*). As a control, we also annotated orthology groups across notothenioid transcriptomes using the OMA pipeline (Altenhoff et al. 2015): 8,697 orthology groups (81%) were consistent with the conservative groups obtained when anchoring the analysis on the stickleback and zebrafish gene sets.

### Amino acid usage in Notothenioid proteins

We sought to examine whether notothenioid proteins exhibited preferential amino-acid usage compared to temperate fish and whether usage differences aligned to the classically accepted psychrophilic modifications in bacteria, such as a decrease in proline, arginine and aromatic residues (Struvay and Feller 2012; Yang et al. 2015). Overall, amino acid compositions inferred from the whole sequence across the 10,728 orthology groups showed no differences between the four notothenioids and the two temperate control fish (stickleback and zebrafish; Figure 4A). We then focused on 45,994 amino acid residue positions where all four notothenioids showed a concordant substitution compared to the amino acid present in temperate fish. These positions were of interest because they likely represented substitutions that occurred during the evolution of notothenioids and were highly unlikely to have resulted from errors in the transcriptome assemblies. At these positions, the Antarctic fish tended to favour serine (S), isoleucine (I) and methionine (M) and disfavour glutamic acid (E) (Figure 4B). The biased usage of serine and methionine seems to be a derived characteristic of notothenioids, because a comparison to more distant human and chicken orthologues yielded similar observations (Supplementary Figure S1). A series of biased amino acid substitutions have taken place in the notothenioids, as shown by the red boxes in Figure 5A. When these substitutions were tested for enrichment, ratio tests showed an imbalance in methionine, isoleucine, lysine (K) and arginine (R) (red boxes, upper most section of Figure 5B). While some ratios were skewed due to low numbers, the most significant bias was the over-representation of positions where leucine in the temperate fish was replaced by a methionine in the notothenioids (lower section of Figure 5B). However, we could not find evidence that these proteins were evolving under positive selection based on comparisons of synonymous vs. non-synonymous substitution rates so the evolutionary impact of those amino-acid substitutions remains unclear (Methods).

**Figure 4.**
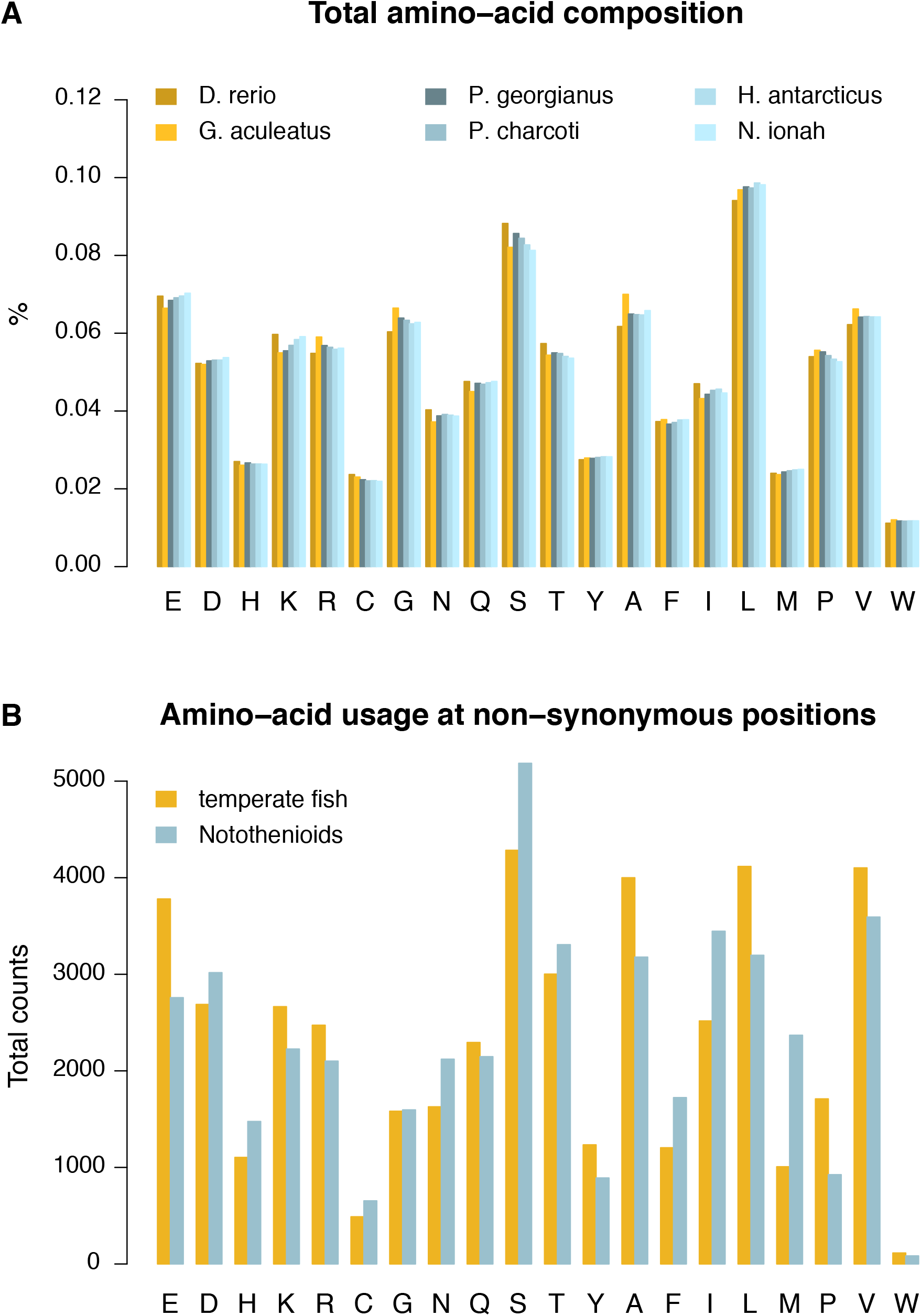
Amino-acid usage in temperate and notothenioid fish proteins. **A.** Total amino-acid composition deduced from the translated sequence of 10,728 orthologous transcripts across species. **B.** Amino-acid usage at 45,994 reliable non-synonymous positions between temperate and notothenioid fish proteins.

**Figure 5.**
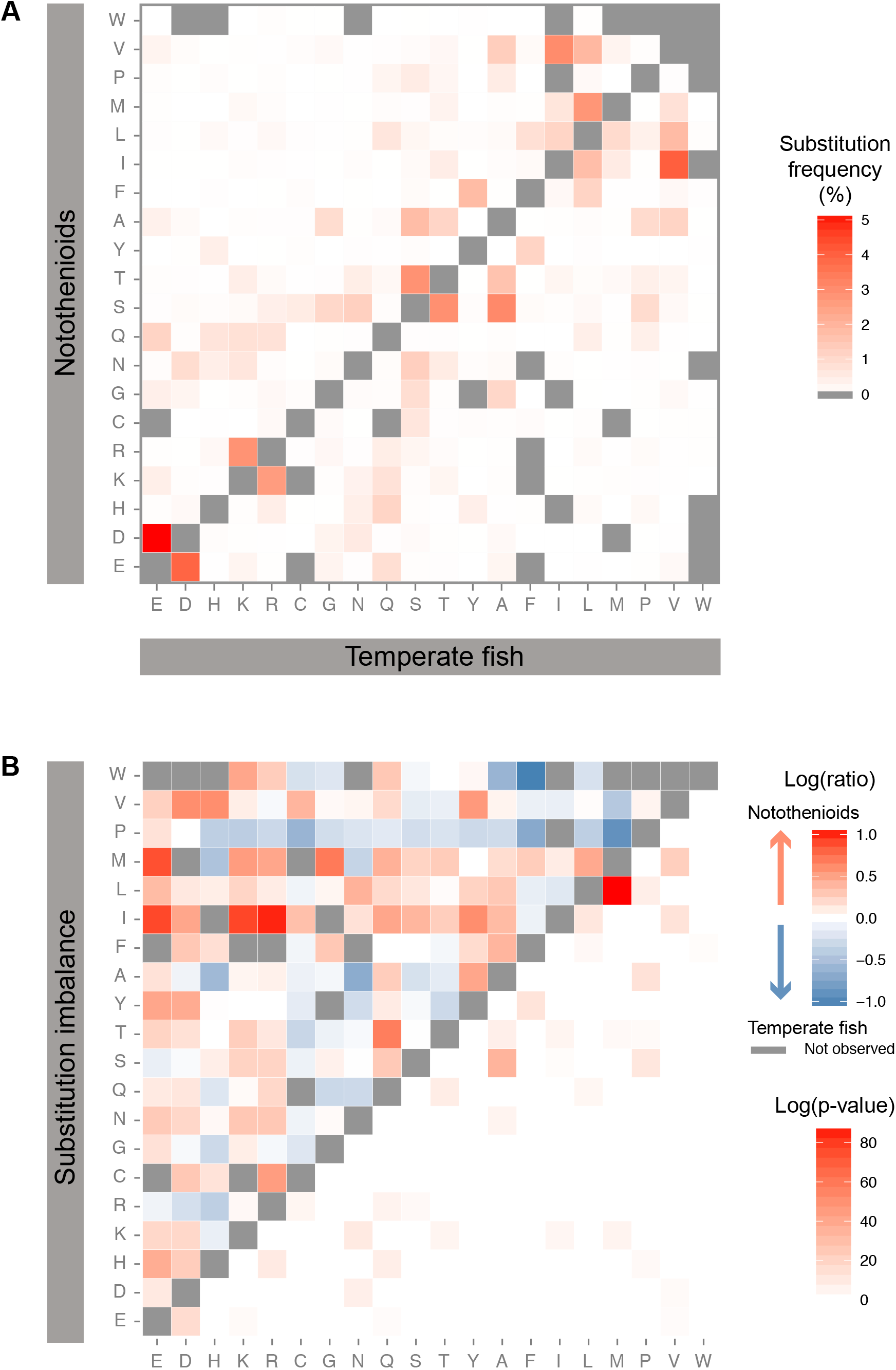
Amino-acid usage imbalance at non-synonymous positions between temperate and notothenioid fish proteins. **A.** Substitution frequencies at 45,994 reliable non-synonymous positions between temperate and notothenioid fish. Percentages are expressed as the fraction across all observed substitutions. Substitutions that were never observed in the curated datasets are shown in grey. **B.** Top left part of the heat map represents the imbalance in the substitution frequencies for any pair of amino-acids between temperate and notothenioid fish. Substitutions favoured in notothenioids are shown in red, while substitutions depleted in notothenioids are shown in blue. Substitutions that are never observed in at least one direction are shown in grey. Bottom right part of the heat map represents the p-value of the proportion test (FDR-corrected, Benjamini-Hochberg adjustment).

### Gene duplications specific to the notothenioids

Of the 10,727 notothenioid orthologous transcripts identified in comparisons with stickleback and zebrafish, 1,628 were identified as potential gene duplications (15% of total) with multiple Antarctic transcripts mapping to a single stickleback transcript. These potential duplicates were manually verified from multiple sequence alignments with their respective stickleback transcripts as mapping to 477 genes, and they were present as multiple copies within at least one Antarctic fish species, rather than multiple non-overlapping transcripts mapping to the same gene. 453 putatively duplicated transcripts were present in at least two Antarctic fish and 46 were present in more than two copies per species. Overall, the potential duplicates mapped to 212 stickleback transcripts (Supplementary Table S2). Multiple sequences mapping to the stickleback transcript ENSGACT00000000161 were removed from further analyses because this transcript putatively encodes a *pol*-like protein within the major histocompatibility locus. This locus contains many repeat elements and thus it was impossible to establish contiguity and correct orthology. Ten other genes that were present in more than two copies in the Antarctic fish were also identified as either being present, absent or in multiple copies in other teleost fish species in a gene-specific manner with no discernible phylogenetic pattern (Supplementary Table S3).

Gene annotation from Ensembl for stickleback (Zerbino et al. 2018) revealed many different putative functions for the stickleback orthologues of Antarctic duplicated transcripts. Analysis of GO terms via the Panther server (Mi et al. 2016) revealed, among notothenioid duplicated genes, an enrichment in the GO-Slim Biological Processes for cell matrix adhesion (6 transcripts, 9.61 fold-enrichment, P = 9.60e^-3^) and cell communication (37 transcripts, 1.81 fold-enrichment, P = 3.65e^-2^), and CCKR (cholecystokinin receptor) signalling was enriched (8 transcripts, 7.99 enrichment, P = 1.40e^-3^) in Panther Pathways analysis (Supplementary Table S4). These observations were confirmed by KEGG analysis, which showed enrichment in transcripts for focal adhesion (9 transcripts P = 2.73e^-3^) and Mapk signalling (6 transcripts, P = 1.94e^-1^). Both Panther and KEGG analyses included the gene *mapk14a* (mitogen-activated protein kinase family member 14a) in the enrichment lists (see below). *mapk14a* is a gene involved in the cellular cascades associated with responses to environmental stress.

Protein-protein interaction analysis using the STRING program and a medium level of confidence revealed a network of 61 proteins centred on the map kinase Mapk14a (Supplementary Figure S2). This network was reduced to 15 interactions, including Mapk14a, when the level of confidence was raised to high (Figure 6, Table 2). This network analysis confirmed the Panther and KEGG analyses, with an emphasis on signalling molecules and transcription factors. The 15 genes in the protein-protein interaction network were only present in one copy, or absent, in other teleosts, but two thirds were present in multiple copies in the evolutionarily distant spotted gar and human genomes (Table 2).

**Table 2.**
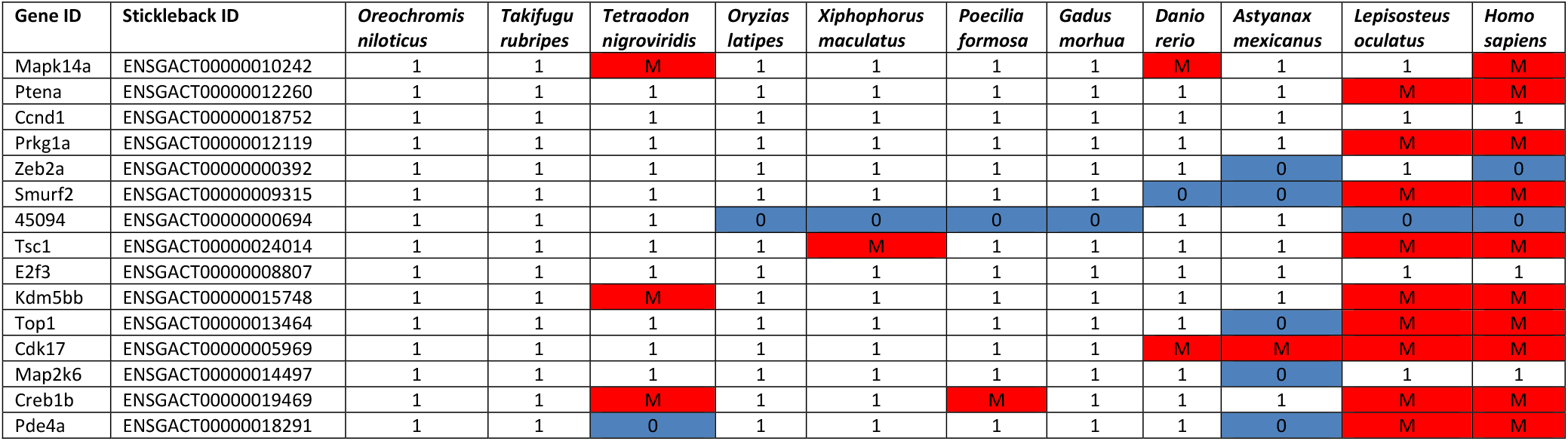
Duplicated genes (two copies or more) in the Antarctic fish in the STRING protein-protein interactions high confidence network. Identification of the number of orthologues in other Ensembl-listed fish species and *Homo sapiens* compared with stickleback, the closely related species to notothenioids in the Ensembl genomes. Key: 0 = absent; 1 = present 1:1 (1 copy in stickleback and 1 copy in the designated species); M = 1: many (1 copy in stickleback and many copies in the designated species).

**Figure 6.**
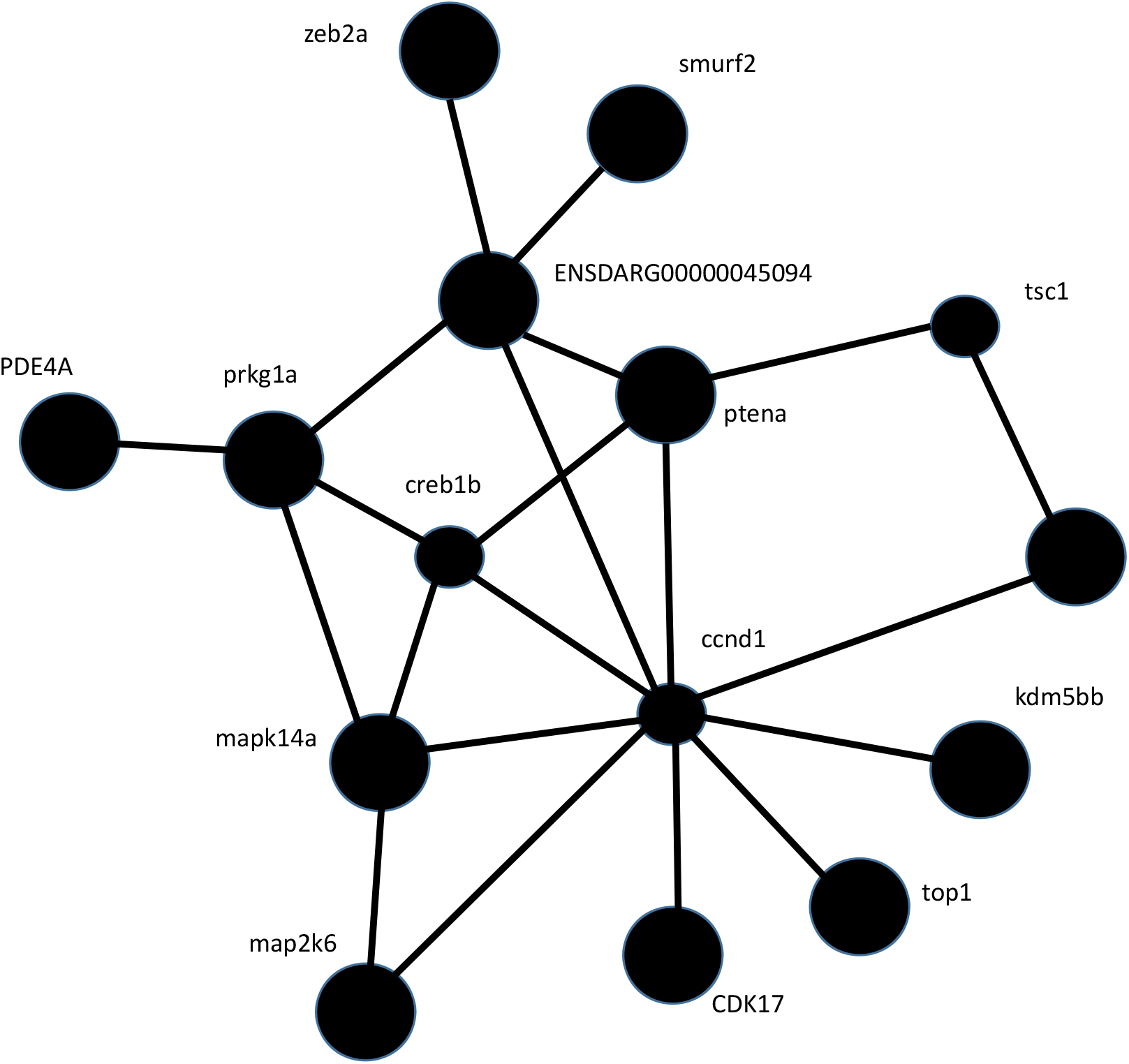
Protein-protein interaction network generated in STRING program at a high confidence level after input of gene identifiers from duplicated genes identified in notothenioids. All genes are involved in either signal/transcriptional transduction or regulation, with the possible exception of ENSDARG00000045094, which has an unknown function and Top1 involved in DNA transcription.

## Discussion

The work presented here provides the first comprehensive comparison of the transcriptomes of multiple Antarctic notothenioids from different taxa with respect to temperate species, thereby negating false positives due to species-specific evolution. The most notable result was the significant over-representation of positions where leucine in the other fish had been replaced by methionine (M, codon AUG) in the notothenioids (Figure 5, bottom heat map). Detailed analyses at concordant, non-synonymous positions between notothenioids and temperate fish showed that Antarctic fish in addition to the significant bias in methionine also favoured serine (S, codons UCN and AGU/C) and isoleucine (I, codons AUU/C/A) over leucine (L, codons CUN), glutamic acid (E, codons GAA/G), lysine (K, codons AAA/G), and arginine (R, codons CGN and AGA/G) (Figures 4 and 5). Similar results for serine and glutamic acid have been reported previously in the Antarctic zoarcid *Pachycara brachycephalum* (Windisch et al. 2012), but the isoleucine and methionine results are novel to this study. The reason behind this significant bias towards methionine remains unclear, especially because both leucine and methionine are non-polar, hydrophobic amino acids and this is not a substitution identified in previous cold adaptation work in bacteria and Archaea (cf. Yang et al. 2015). For such a change to occur, in most instances two nucleotide mutations would be required in the triplet code to convert leucine (CUU, CUC, CUA, CUG, UUA or UUG) into methionine (AUG), which would be a rare event. Methionine residues are rare in vertebrate proteins (<2%) and are normally quite conserved. There are several possible hypotheses as to why this bias in methionines evolved including increasing protein stability, alternative start sites and redox sensors.

Although the incorporation of methionine as opposed to a branched amino acid would give increased flexibility in the core (Pucci and Rooman 2017), this situation can be energetically costly and its effectiveness would obviously depend on the position of the methionine:leucine substitution. However, the fact that the more bulky isoleucine is preferred over the similarly beta-branched valine does not suggest that there has been pressure to reduce packing in the core of cold adapted proteins. Mutation studies in a bacterial membrane protein revealed that replacement of methionines by leucines reduced stability and affected protein-lipid interactions (Chaturvedi and Mahalakshimi 2013). Hence in the bacterial membrane protein, the methionines were external, but still increased the stability of the protein at 4°C.

An alternative hypothesis is that these “extra” methionines may act as alternative start sites for certain genes, but a more plausible explanation could be that they are involved in redox regulation. The freezing, oxygen-rich waters of the Southern Ocean promote the formation of reactive oxygen species (ROS) and would be expected to lead to enhanced ROS damage of DNA and membrane lipid peroxidation in polar species (Abele and Puntarulo 2004). To promote the protection of the genome, particularly in long-lived species like notothenioid fish (Buttemer et al. 2010), the expectation would be the replacement of methionines with leucines, or similar amino acids because the sulphur-containing amino acids (methionine and cysteine) are targets for ROS. Indeed a reduction in methionines has been identified in psychrophilic prokaryotes when compared with thermophilic species (Yang et al. 2015). In mammals, however, these sulphur-containing amino acids have been shown to act as critical antioxidant regulators (Stadtman et al. 2002). Both methionine and cysteine can undergo reversible modifications to modulate physiological protein functions and acting as molecular redox switches (Pajares et al. 2015)., As supporting evidence for this hypothesis, studies on two Antarctic bivalve molluscs (*Aequiyoldia eightsi*, previously *Yoldia eightsi*) and *Adamussium colbecki*) have shown trade-offs between the thermal tolerance and antioxidation potential of the animal (Regoli et al. 1997; Abele et al. 2001). In both instances, the Antarctic antioxidant enzymes catalase and superoxide dismutase (SOD) are highly efficient at 0°C, more so than the orthologous enzymes in closely related temperate relatives, but these Antarctic enzymes lose their activity with even small increases in temperature (Regoli et al. 1997; Abele et al. 2001). The resulting hypothesis was that at least some of the enzymes involved in the Antarctic bivalve antioxidant system have been fine-tuned to work at freezing temperatures. However, these particular enzymes cannot maintain their 3-dimensional integrity when temperatures are increased and thus these enzymes cease to function when conditions warm (Regoli et al. 1997; Abele et al. 2001). Recent work in Antarctic fish examined antioxidant activity in three species related to warming (Enzor and Place 2014). Whilst all three species elevated total SOD and catalase activity to short-term increases in water temperatures to 4°C, this was not sustained long-term. In addition at the end of the two month experiment levels of cellular damage, as measured by protein carbonyls, remained slightly above those of control animals, indicating incomplete acclimation to 4°C after 56 days and potential trade-offs at the cellular level to chronic oxidative damage or damage from other sources (Enzor and Place 2014).

The results described here for notothenioid fish differ from other studies on psychrophiles. In prokaryotic studies specific amino acid substitutions have been associated with cold adaptation, including a reduced use of proline (P), arginine and acidic amino acids in bacteria, a lower content of hydrophobic amino acids (leucine in particular) in the Archaea and a higher content of non-charged polar amino acids, especially glycine (G) and threonine (T) in bacteria (Saunders et al. 2003, Ayala-del-Rio et al. 2010; Zhao et al. 2010; Yang et al. 2015). These changes result in increased flexibility when compared with mesophilic proteins, especially of the active site, which is deemed essential for efficient enzyme functioning in the cold (Struvay & Feller 2012). However, it is still not possible to predict *a priori* with any great accuracy how particular amino acid substitutions will affect protein activity at different temperatures (Fields et al. 2015). Although increasingly sophisticated algorithms are being developed to predict protein activity in response to temperature changes and protein stability following mutations, accurate prediction of reaction rates of enzymes across a temperature gradient remains difficult, not least because all interactions are context-dependent (Fields et al. 2015; Pucci and Rooman 2017). This lack of ability to predict enzymatic activity related to temperature is due to an intricate combination of different intrinsic and extrinsic factors that frequently differ according to the protein or protein family and which are difficult to disentangle (Struvay and Feller 2012; Fields et al. 2015; Pucci and Rooman 2017). In many cases, alteration in enzyme performance can result from a single amino acid substitution which can be demonstrated only by functional analyses (Fields et al. 2015). The situation is further complicated by the fact that different proteomic amino acid compositional motifs have been demonstrated between archaea, eubacteria and eukaryotes (Pe’er et al. 2003), which may partly explain why Antarctic fish display different amino acid substitutions than those identified in bacteria and Archaea.

It is now clear from prokaryotic analyses that, in terms of amino acid composition, low temperature adaptation is not necessarily the opposite of high temperature adaptation with kinetic stabilisation against cold denaturation suggested as a cold adaptation mechanism (Romero-Romero et al. 2011; Yang et al. 2015). However, with the vast majority of data underpinning these observations originating from single celled organisms (especially bacteria) (cf. Methe et al. 2005; Struvay and Feller 2012; Yang et al. 2015; Pucci and Rooman 2017) it may well be that the situation is more complex in psychrophilic multicellular organisms, in which there is a relative paucity of both sequence data and functional analyses.

Only a handful of protein adaptations have been functionally described in Metazoa living at sub-zero temperatures. Published studies of some Metazoa do, however, include some of the adaptations detailed above leading to the evolution of proteins with higher maximum activity and lower temperature of maximum stability, e.g. lactate dehydrogenase, tubulins, L-glutamate dehydrogenase, the transmembrane protein *Sec61*, and the TCP chaperone complex in Antarctic fish (Fields & Somero 1998; Detrich et al. 2000; Ciardiello et al. 2000; Römisch et al. 2003; Pucciarelli et al. 2006; Cuellar et al. 2014) as well as the evolution of novel variants such as nothepsins in the aspartate protease family (De Luca et al. 2009). For some other genes (e.g., tubulins and pepsins) relative protein production levels in notothenioids are similar to those in temperate species due to gene duplication events in Antarctic species, where two genes produce similar amounts of protein as one gene in temperate relatives (Parker and Detrich 1998; Carginale et al. 2004). In this study, a number of potential gene duplications were identified when orthology assignments were carried out. Manual verification confirmed that 453 transcripts were duplicated in the notothenioids within our orthologous dataset (Supplementary Tables S2 and S3). Enrichment and network analyses demonstrated the importance of focal adhesion and cell signalling, especially the MAPK pathway in these duplicated genes (Table 2, Figure 6, Supplementary Figure S2 and Supplementary Table S4). Network analysis showed a single node clustered around Mapk14a. This serine kinase is an essential component of the MAP kinase signal transduction pathway and is one of four p38 MAPKs playing an important role in the cellular cascades evoked by environmental stress (Uren Webster and Santos 2015), a phenomenon well documented in marine species and clearly important in Antarctic species that have a requirement to combat high levels of ROS (e.g. Notch et al. 2012).

Gene duplication within the notothenioids has previously been demonstrated in other analyses of notothenioid transcriptomes (cf. Chen et al. 2008, Coppe et al. 2012, Windisch et al. 2012). Each study largely showed duplication of different biochemical pathways, such as proteins associated with mitochondrial localisation, mitochondrial function and biogenesis and ROS genes (Coppe et al. 2012), immune function and fatty acid catabolism (Windisch et al. 2012) and the electron transport chain, transcriptional regulation, protein binding and the ubiquitin conjugated pathway (Chen et al. 2008; Shin et al. 2012; Bilyk and Cheng 2013). The most comprehensive analysis to date was in the *Dissostichus mawsoni* EST study (Chen et al. 2008), where comparative genome hybridization was used to identify around 200 genes augmented in Antarctic species. Of direct relevance to this study was the identification of numerous “stress” genes including ten involved in the Ras/MAPK pathway, elements of which were also shown to be enriched in our data presented here.

To date, the vast majority of analyses of Antarctic fish genes, including the one described here, are largely constrained by the fact that the interrogated datasets are all transcriptomes. Many transcriptome studies focus on either single species, or a limited number of tissues and are therefore, reliant on the duplicated genes being sufficiently represented to enable pathway enrichment detection by either PANTHER or KEGG within these datasets. Additionally, many of these studies compare a single species of Antarctic fish with a single temperate relative and therefore cannot avoid any potential lineage-specific, species-specific or tissue-specific biases. Analysis of the amino acid substitutions in several candidate proteins emphasised the need to study such changes across multiple species. Three selected proteins studied here (calmodulin, neuroglobin and SOD1 (Supplementary Information S1)), showed more species-specific amino acid substitutions within notothenioids than between notothenioids and other fish species. In addition, the positions and types of proposed “notothenioid-specific” changes were invariably in the same coding sites that showed high variability among other fish species, with substitutions at the same position of up to six different amino acid residues with many different properties (Supplementary Information S1). Thus, the question of how extensive gene duplication events are in the notothenioids can only be answered with whole genome sequencing of multiple species. In addition, the extent to which these species use the different strategies of either amino acid substitutions to increase protein flexibility and/or gene duplication to increase protein quantity in the cold is unknown. The identification of such changes in proteins selected so far for functional studies may be serendipitous (e.g., most tubulins are notoriously cold-sensitive) and not follow the general rule.

The global analyses performed here provided little evidence for mass alteration of amino acid composition of Antarctic fish proteins (with the exception of methionines) that might have been predicted to enhance protein folding in the cold and reduce cold denaturation of protein, yet these fish are stenothermal (Somero and Devries 1967; Bilyk and Devries 2011). There is evidence that protein un-folding and accumulation of ubiquitinated proteins may be a significant problem in Antarctic species (Peck 2016, 2018). Analyses have shown that Antarctic fish contain higher levels of ubiquitin-tagged proteins than closely related temperate species (Place and Hofmann 2005; Todgham et al. 2007, 2017). In addition, Antarctic fish permanently express the inducible form of the 70kDa heat shock protein (HSP70), a molecular chaperone that targets denatured and unfolded proteins (Hofmann et al. 2000; Place et al. 2004; Clark et al. 2008). However, recent evidence suggests that the activity of the proteasome apparatus may be temperature-compensated in at least some tissues of notothenioid species (measured in *Trematomus bernacchii* and *Pagothenia borchgrevinki*) (Todgham et al. 2017). Gene expression analyses of the Ub-proteosome pathways suggests that this may be due to higher concentrations of proteasomes in the cell, although the catalytic efficiency of the 20S proteasome has yet to be evaluated (Todgham et al. 2017). Clearly this compensation is not sufficient to avoid the accumulation of unfolded proteins and the mechanism leading to this accumulation remains unknown. Thus, at the protein amino acid sequence level, these fish appear to be poorly adapted to the cold. This finding may explain the high levels of ubiquitination found in these species, but would not explain the stenothermality of these fish (Somero and Devries 1967; Bilyk and Devries 2011). The additional question remains about the function of the increase in the number of methionines. Does the preferential assimilation of this rare and specialised amino acid into the genome increase protein stability in critical proteins or to act as a redox sensor? These cellular results contrast with the ecological situation, whereby the notothenioids are highly successful. Future vulnerabilities will almost certainly be the result of complex interactions including the cellular level constraints detailed above and physiological and ecological characteristics such as very low metabolic rates, low energy lifestyles and limited food supply (Peck 2016, 2018).

## Summary

Even in bacteria, it is difficult using *in silico* approaches to define changes that enable a protein to function efficiently in the cold. It is also suggested that adaptation to the psychrophilic lifestyle is more the result of a suite of synergistic rather than unique changes (Methe et al. 2005). To date, because bioinformatic approaches still have limited success in predicting function from sequence, hence there is a need to move towards high through-put testing of psychrophilic metazoan proteins to progress our understanding beyond that of a few isolated proteins and to develop an overview of genome-wide cold adaptations. The new era of long read sequencing will facilitate the generation of reference-quality genomes, and the notothenioids are currently one of the specialist groups targeted by the Sanger Institute as part of their contribution to the international Vertebrate Genome Project. These genomes will provide an invaluable resource for functional studies, which are clearly essential for understanding the subtle amino acid substitutions identified in this study, particularly the role arising from the significant increase in methionines. Given evidence accumulated to date, it is entirely possible that the Antarctic marine fauna are “clinging on” to life in their environment with only a selected number of biochemical pathways modified to work efficiently below 0°C (Clark et al. 2017). In a warming world, it may be this relatively small number of cold-adapted proteins that are responsible for the vulnerability of these stenothermal species with considerable impacts for ecosystem functioning in the future.

## Materials and Methods

*Material: Harpagifer antarcticus* (Nybelin, 1947) were collected by divers from a depth of 12 m from Ryder Bay (67°34’07” S, 68°07’30” W), close to the Rothera research station run by the British Antarctic Survey on the Antarctic Peninsula. Seven tissues were dissected and flash frozen in liquid nitrogen for subsequent extraction of RNA. Specimens of the dragonfish *Parachaenichthys charcoti* (Vaillant, 1906) and of the South Georgia icefish *Pseudochaenichthys georgianus* (Norman, 1937) were collected by bottom trawls deployed from the *ARSV Laurence M. Gould* south of Low Island (Antarctic Specially Protected Area No. 152, Western Bransfield Strait) or west of Brabant Island (Antarctic Specially Protected Area 153, Eastern Dallmann Bay) in the Palmer Archipelago (March-June, 2012 and 2013). Fourteen and ten tissues were dissected respectively from each species and flash frozen in liquid nitrogen for subsequent extraction of RNA. A single specimen of the icefish *Neopagetopsis ionah* (Nybelin, 1947) was captured by bottom trawling in Andvord Bay (May, 2012) near Palmer Station. Only spleen tissue was available from this species (Supplementary Table S1).

### Transcriptome sequencing and assembly

Total RNA was extracted from 58 samples across the four species using TRIsure (Bioline) according to manufacturer’s instructions and purified on RNeasy columns (Qiagen). Quantity and quality were checked on an Agilent TapeStation 2200. The concentrated RNA samples were submitted to the EMBL GeneCore facility (Heidelberg, Germany) and used to generate barcoded normalized cDNA libraries with an average fragment length of ~130 bp. The libraries were multiplexed and sequenced over four lanes on a HiSeq2000 platform (Illumina, San Diego, USA) using 100 bp paired-end reads. Read quality was assessed using FastQC (http://www.bioinformatics.babraham.ac.uk/projects/fastqc/). For each of the four species, raw sequencing reads from the different tissues (Supplementary Table S1) were pooled for full transcriptome assembly performed with Trinity (Grabherr et al. 2011), using Trimmomatic (Bolger et al. 2014) to pre-process and clip low-quality reads and remaining adapter sequences, and default parameters otherwise. Sequencing read files are available from ArrayExpress (accession number E-MTAB-6759) and assembled transcriptomes are available from BioStudies (accession number S-BSST132).

### Transcriptome quality control

Reads used for the *de novo* assemblies were remapped to the assembled transcripts using Bowtie (Langmead et al. 2009) with the parameters implemented in the RSEM pipeline (Li and Dewey 2011). Transcriptome completeness was estimated with BUSCO using the *Eukaryota, Metazoa, Vertebrata* and *Actinopterygii* reference gene sets (Simão et al. 2015).

### Orthology assignments

For sequence evolution analyses, transcripts were additionally assigned to orthologous gene families based on three-way orthologous relationships with stickleback (*G. aculeatus*) and zebrafish (*D. rerio*). Curated, non-redundant cDNA sets for stickleback and zebrafish were downloaded from Ensembl v81 (Cunningham et al. 2015) and compared to the four notothenioid transcriptome assemblies independently using BLASTX to identify putative orthologs (–max_target_seqs 1, –evalue 1e-40) (Altschul et al. 1990). Family assignation was considered robust when the stickleback and zebrafish transcripts separately identified by the BLASTX analysis of the four notothenioid transcriptome assemblies themselves corresponded to a pair of orthologous genes according to the Ensembl Compara gene trees (Vilella et al. 2009). Transcripts that could not be assigned an orthologue in either of the two species, or resulted in a discrepancy between stickleback and zebrafish (i.e. the best BLAST hits were identified as paralogous in the Ensembl Compara gene trees, or otherwise non-orthologous genes) were not used for sequence evolution analyses. The OMA pipeline was also run with default parameters on the four notothenioid transcriptomes to independently identify orthologues across notothenioids (Altenhoff et al. 2015). Orthology groups were considered as consistent when a group of orthologous notothenioid transcripts identified by one method fully overlapped the group identified by the other and were kept for analyses.

### Sequence alignments

Once transcripts were assigned to an orthologous gene family, the Ensembl API was used to retrieve the orthologous cDNA sequences in all 11 fish available in the database ((*Danio rerio* (Zebrafish), *Gasterosteus aculeatus* (Stickleback), *Astyanax mexicanus* (Cavefish), *Gadus morhua* (Cod), *Oreochromis niloticus* (Tilapia), *Oryzias latipes* (Medaka), *Xiphophorus maculatus* (Platyfish), *Poecilia formosa* (Amazon molly), *Takifugu rubripes* (Fugu or the Japanese pufferfish), *Tetraodon nigroviridis* (Tetraodon or the green spotted pufferfish), *Leipisteus oculatus* (Spotted gar)), as well as human (*Homo sapiens*) and chicken (*Gallus gallus*) as outgroups. The orthologous transcripts were translated to protein sequences and the protein sequences aligned using T-Coffee with default parameters (Notredame et al. 2000). The alignments were then back-translated to the transcript sequences using the “backtrans” utility from TreeBeST (Vilella et al. 2009). Low-quality blocks in the sequence alignments were removed using GBlocks (–t=c, –b5=h) (Castresana 2000). Gene trees reconciled with the species phylogeny were built using TreeBeST v.1.9.2 with default parameters.

### Analyses of amino acid composition and substitutions

Global amino-acid composition was computed across species on translated orthologous transcripts to ensure that missing or redundant transcripts in the notothenioid transcriptomes would not skew the comparisons. High-quality blocks were extracted from the protein alignments using the “seq_reformat” utility in T-Coffee (minimum column score of 8). The filtered alignments were then parsed using a custom Python script to identify sites where notothenioids exhibit the same amino-acid substitution compared to reference temperate fish (stickleback and zebrafish), i.e. a presumably fixed substitution that occurred in the ancestral notothenioid branch.

### Positive selection scans

Codeml (Yang 2007) was run on the sequence alignments and reconciled gene trees (described above) using the *ETE3* Python package (Heurta-Cepas et al. 2016) to detect proteins with signs of either positive selection or constraint relaxation specifically in Notothenioids. Synonymous and non-synonymous substitution rates were estimated using branch models with the ancestral Notothenioidei branch as the foreground and other branches in the tree as the background (null model ‘M0’, free evolution of the foreground ‘b_free’ and neutral evolution of the foreground ‘b_neut’). The two non-null models were compared to the null model for each tree using a likelihood ratio test.

### Determination of duplicated Notothenioid genes

Potential Antarctic-specific gene duplications were identified when multiple Antarctic sequences mapped to a single stickleback transcript. These alignments were manually verified to determine whether these multiple mappings were the result of smaller Antarctic transcripts mapping to different parts of a longer stickleback transcript, or whether there was sufficient overlap in the Antarctic sequences to assign them as duplicated genes. Extraction of the number of orthologues in other fish species and *H. sapiens* was carried out via Ensembl. *G. aculeatus* assigned gene names for these duplicated transcripts were used to perform Gene Ontology (GO) analysis via the Panther server (Mi et al. 2016) and identify KEGG pathways and protein-protein interactions via STRING v10 (Szklarczyk et al. 2015).

## Acknowledgements and funding information

This work was supported by an NERC/Cambridge University Innovation Award (MSC, LSP, JC, AN, PF and CB); by U.S. National Science Foundation (ANT-1247510 and PLR-1444167 to HWD; PLR-1543383 to JHP, TD & HWD); by the Wellcome Trust (WT108749/Z/15/Z to PF); the European Molecular Biology Laboratory (CB and PF) and NERC core funding to the British Antarctic Survey (MSC and LSP). HWD, JHP and TD gratefully acknowledge the logistic support provided to their Antarctic field research program, performed at Palmer Station and on the seas of the Palmer Archipelago, by the staff of the Division of Polar Programs of the National Science Foundation, by the personnel of the Antarctic Support Contractor, and by the captains and crews of the *ARSV Laurence M. Gould*. MSC, LSP and JC gratefully acknowledge the help of Adrian Nickson (AN) in the early stages of this project, including obtaining the Innovation Award. The authors would like to thank all members of the Rothera Dive Team for providing *Harpagifer antarcticus* samples and Jamie Oliver at BAS for help with illustrations and Laura Gerrish at BAS for the map. Overall BAS diving support was provided by the NERC National Facility for Scientific Diving at Oban. The EMBL GeneCore is acknowledged for sequence data production. This is contribution #XXX from the Marine Science Center at Northeastern University. Sequencing read files are available from ArrayExpress (accession number E-MTAB-6759) and assembled transcriptomes are available from BioStudies (accession number S-BSST132).

## References

Abele D, Puntarulo S. 2004. Formation of reactive species and induction of antioxidant defence systems in polar and temperate marine invertebrates and fish. Comp Biochem Physiol A Mol Integr Physiol. 138:405–415.

Abele D, Tesch C, Wencke P, Portner HO. 2001. How does oxidative stress relate to thermal tolerance in the Antarctic bivalve *Yoldia eightsi*? Antarct Sci. 13:111–118.

Abram NJ, Mulvaney R, Wolff E, Triest J, Kipfstuhl S, Trusel LD, Vimeux F, Fleet L, Arrowsmith C. 2013. Acceleration of snow melt in an Antarctic Peninsula ice core during the twentieth century. Nat Geosci. 6:404–411.

Altenhoff AM, Skunca N, Glover N, Train CM, Sueki A, Pilizota I, Gori K, Tomiczek B, Muller S, Redestig H, et al. 2015. The OMA orthology database in 2015: function predictions, better plant support, synteny view and other improvements. Nucl Acids Res. 43:D240–D249.

Altschul SF, Gish W, Miller W, Myers EW, Lipman DJ. 1990. Basic local alignment search tool. J Mol Biol. 215:403–410.

Ayala-del-Rio HL, Chain PS, Grzymski JJ, Ponder MA, Ivanova N, Bergholz PW, Di Bartolo G, Hauser L, Land M, Bakermans C, et al. 2010. The genome sequence of *Psychrobacter arcticus* 273-4, a psychroactive Siberian permafrost bacterium, reveals mechanisms for adaptation to low-temperature growth. Appl Environ Microbiol. 76:2304–2312.

Betancur-R R, Broughton RE, Wiley EO, Carpenter K, Lopez JA, Li C, Holcroft NI, Arcila D, Sanciangco M, Cureton Ii JC, et al. 2013. The tree of life and a new classification of bony fishes. PLoS Curr. 5:2013 April 18doi: 10.1371/currents.tol.53ba26640df0ccaee75bb165c8c26288

Billger M, Wallin M, Williams RC, Detrich HW. 1994. Dynamic instability of microtubules from cold-living fishes. Cell Motil Cytoskeleton 28:327–332.

Bilyk KT, Cheng CHC. 2013. Model of gene expression in extreme cold - reference transcriptome for the high-Antarctic cryopelagic notothenioid fish *Pagothenia borchgrevinki*. BMC Genomics 14:634

Bilyk KT, Cheng CHC. 2014. RNA-seq analyses of cellular responses to elevated body temperature in the high Antarctic cryopelagic nototheniid fish Pagothenia borchgrevinki. Mar Genomics 18:163–171.

Bilyk KT, Devries AL. 2011. Heat tolerance and its plasticity in Antarctic fishes. Comp Biochem Physiol A Mol Integr Physiol. 158:382–390.

Bolger AM, Lohse M, Usadel B. 2014. Trimmomatic: a flexible trimmer for Illumina sequence data. Bioinform. 30:2114–2120.

Buttemer WA, Abele D, Costantini D. 2010. From bivalves to birds: oxidative stress and longevity. Funct Ecol. 24:971–983.

Capasso C, Lees WE, Capasso A, Scudiero R, Carginale V, Kille P, Kay J, Parisi E. 1999. Cathepsin D from the liver of the Antarctic icefish *Chionodraco hamatus* exhibits unusual activity and stability at high temperatures. Biochim Biophys Acta-Protein Structure and Molecular Enzymology 1431:64–73.

Carginale V, Trinchella F, Capasso C, Scudiero R, Parisi E. 2004. Gene amplification and cold adaptation of pepsin in Antarctic fish. A possible strategy for food digestion at low temperature. Gene 336:195–205.

Carpenter JH. 1966. New measurements of oxygen solubility in pure and natural water. Limnol. Oceanogr. 11:264–277.

Castresana J. 2000. Selection of conserved blocks from multiple alignments for their use in phylogenetic analysis. Mol Biol Evol. 17:540–552.

Chaturvedi D, Mahalakshmi R. 2013. Methionine mutations of outer membrane protein X influence structural stability and beta-barrel unfolding. PLoS One 8: e79351

Chen LB, Devries AL, Cheng CHC. 1997. Evolution of antifreeze glycoprotein gene from a trypsinogen gene in Antarctic notothenioid fish. Proc Natl Acad Sci USA 94:3811–3816.

Chen ZZ, Cheng CHC, Zhang JF, Cao LX, Chen L, Zhou LH, Jin YD, Ye H, Deng C, Dai ZH, et al. 2008. Transcrintomic and genomic evolution under constant cold in Antarctic notothenioid fish. Proc Natl Acad Sci USA 105:12944–12949.

Ciardiello MA, Camardella L, Carratore V, di Prisco G. 2000. L-glutamate dehydrogenase from the Antarctic fish Chaenocephalus aceratus - Primary structure, function and thermodynamic characterisation: relationship with cold adaptation. Biochim Biophys Acta - Protein Structure and Molecular Enzymology 1543:11–23.

Clark MS, Fraser KPP, Burns G, Peck LS. 2008. The HSP70 heat shock response in the Antarctic fish *Harpagifer antarcticus*. Polar Biol. 31:171–180.

Clark MS, Sommer U, Sihra JK, Thorne MAS, Morley SA, King M, Viant MR, Peck LS. 2017. Biodiversity in marine invertebrate responses to acute warming revealed by a comparative multi-omics approach. Glob Change Biol. 23:318–330.

Coppe A, Agostini C, Marino IAM, Zane L, Bargelloni L, Bortoluzzi S, Patarnello T. 2013. Genome evolution in the cold: Antarctic icefish muscle transcriptome reveals selective duplications increasing mitochondrial function. Genome Biol Evol. 5:45–60.

Cuellar J, Yebenes H, Parker SK, Carranza G, Serna M, Valpuesta JM, Zabala JC, Detrich HW. 2014. Assisted protein folding at low temperature: evolutionary adaptation of the Antarctic fish chaperonin CCT and its client proteins. Biol Open 3:261–270.

Cunningham F, Amode MR, Barrell D, Beal K, Billis K, Brent S, Carvalho-Silva D, Clapham P, Coates G, Fitzgerald S, et al. 2015. Ensembl 2015. Nucl Acids Res. 43:D662–D669.

De Luca V, Maria G, De Mauro G, Catara G, Carginale V, Ruggiero G, Capasso A, Parisi E, Brier S, Engen JR, et al. 2009. Aspartic proteinases in Antarctic fish. Mar Genomics 2:1–10.

Detrich HW, III, Johnson KA, Marschese-Ragona SP. 1989. Polymerisation of Antarctic fish tubulins at low temperatures energetic aspects. Biochem. 28:10085–10093.

Detrich HW, Parker SK, Williams RC, Nogales E, Downing KH. 2000. Cold adaptation of microtubule assembly and dynamics - Structural interpretation of primary sequence changes present in the alpha- and beta-tubulins of Antarctic fishes. J Biol Chem. 275:37038–37047.

Devries AL. 1971. Glycoproteins as biological antifreeze agents in Antarctic fishes. Science 172:1152–1155.

di Prisco G, Cocca E, Parker SK, Detrich HW. 2002. Tracking the evolutionary loss of hemoglobin expression by the white-blooded Antarctic icefishes. Gene 295:185–191.

Dunton K. 1992. Arctic biogeography – the paradox of the marine benthic fauna and flora. Trends Ecol Evol. 7:183–189.

Eastman JT, Eakin RR. 2000. An updated species list for notothenioid fish (Perciformes; Notothenioidei), with comments on Antarctic species. Arch Fish Mar Sci. 48:11–20.

Enzor LA, Place SP. 2014. Is warmer better? Decreased oxidative damage in notothenioid fish after long-term acclimation to multiple stressors. J Exp Biol. 217:3301–3310.

Fields PA, Dong YW, Meng XL, Somero GN. 2015. Adaptations of protein structure and function to temperature: there is more than one way to ‘skin a cat’. J Exp Biol. 218:1801–1811.

Fields PA, Somero GN. 1998. Hot spots in cold adaptation: Localized increases in conformational flexibility in lactate dehydrogenase A(4) orthologs of Antarctic notothenioid fishes. Proc Natl Acad Sci USA 95:11476–11481.

Gerdol M, Buonocore F, Scapigliati G, Pallavicini A. 2015. Analysis and characterization of the head kidney transcriptome from the Antarctic fish *Trematomus bernacchii* (Teleostea, Notothenioidea): A source for immune relevant genes. Mar Genomics 20:13–15.

Grabherr MG, Haas BJ, Yassour M, Levin JZ, Thompson DA, Amit I, Adiconis X, Fan L, Raychowdhury R, Zeng QD, et al. 2011. Full-length transcriptome assembly from RNA-Seq data without a reference genome. Nature Biotechnol. 29:644–U130.

Haas BJ, Papanicolaou A, Yassour M, Grabherr M, Blood PD, Bowden J, Couger MB, Eccles D, Li B, Lieber M, et al. 2013. *De novo* transcript sequence reconstruction from RNA-seq using the Trinity platform for reference generation and analysis. Nat Protoc. 8:1494–1512.

Herrero J, Muffato M, Beal K, Fitzgerald S, Gordon L, Pignatelli M, Vilella AJ, Searle SMJ, Amode R, Brent S, et al. 2016. Ensembl comparative genomics resources. Database (2016) Vol. 2016: article ID bav096; doi: 10.1093/database/bav096

Himes RH, Detrich HW. 1989. Dynamics of Antarctic fish microtubules at low temperatures. Biochem. 28:5089–5095.

Hofmann GE, Buckley BA, Airaksinen S, Keen JE, Somero GN. 2000. Heat-shock protein expression is absent in the Antarctic fish *Trematomus bernacchii* (family Nototheniidae). J Exp Biol. 203:2331–2339.

Huerta-Cepas J, Serra F, Bork P. 2016. ETE 3: Reconstruction, analysis, and visualization of phylogenomic data. Mol. Biol. Evol. 33:1635–1638.

Hunt GL, Drinkwater KF, Arrigo K, Berge J, Daly KL, Danielson S, Daase M, Hop H, Isla E, Karnovsky N, et al. 2016. Advection in polar and sub-polar environments: Impacts on high latitude marine ecosystems. Prog Oceanogr.149:40–81.

Huth TJ, Place SP. 2013. De novo assembly and characterization of tissue specific transcriptomes in the emerald notothen, *Trematomus bernacchii*. BMC Genomics 14:805

Johnston IA, Fernandez DA, Calvo J, Vieira VLA, North AW, Abercromby M, Garland T. 2003. Reduction in muscle fibre number during the adaptive radiation of notothenioid fishes: a phylogenetic perspective. J Exp Biol. 206:2595–2609.

Langmead, Trapnell C, Pop M, Salzberg SL (2009) Ultrafast and memory-efficient alignment of short DNA sequences to the human genome. Genome Biol. 10:R25.

Lecointre G, Ameziane N, Boisselier MC, Bonillo C, Busson F, Causse R, Chenuil A, Couloux A, Coutanceau JP, Cruaud C, et al. 2013. Is the species flock concept operational? The Antarctic shelf case. PLoS One 8: e68787.

Li B, Dewey CN. 2011. RSEM: accurate transcript quantification from RNA-Seq data with or without a reference genome. BMC Bioinform. 12:323.

Maldonado A, Bohoyo F, Galindo-Zaldivar J, Hernandez-Molina FJ, Lobo FJ, Lodolo E, Martos YM, Perez LF, Schreider AA, Somoza L. 2014. A model of oceanic development by ridge jumping: Opening of the Scotia Sea. Glob Planetary Change 123:152–173.

Methe BA, Nelson KE, Deming JW, Momen B, Melamud E, Zhang XJ, Moult J, Madupu R, Nelson WC, Dodson RJ, et al. 2005. The psychrophilic lifestyle as revealed by the genome sequence of *Colwellia psychrerythraea* 34H through genomic and proteomic analyses. Proc Natl Acad Sci USA 102:10913–10918.

Mi HY, Poudel S, Muruganujan A, Casagrande JT, Thomas PD. 2016. PANTHER version 10: expanded protein families and functions, and analysis tools. Nucl Acids Res. 44:D336–D342.

Moylan TJ, Sidell BD. 2000. Concentrations of myoglobin and myoglobin mRNA in heart ventricles from Antarctic fishes. J Exp Biol. 203:1277–1286.

Near TJ, Dornburg A, Kuhn KL, Eastman JT, Pennington JN, Patarnello T, Zane L, Fernandez DA, Jones CD. 2012. Ancient climate change, antifreeze, and the evolutionary diversification of Antarctic fishes. Proc Natl Acad Sci USA 109:3434–3439.

Nicolas JP, Bromwich DH. 2014. New reconstruction of Antarctic near-surface temperatures: multidecadal trends and reliability of global reanalyses. J Clim. 27:8070–8093.

Notch EG, Chapline C, Flynn E, Lameyer T, Lowell A, Sato D, Shaw JR, Stanton BA. 2012. Mitogen activated protein kinase 14-1 regulates serum glucocorticoid kinase 1 during seawater acclimation in Atlantic killifish, Fundulus heteroclitus. Comp Biochem Physiol A Mol Integr Physiol. 162:443–448.

Notredame C, Higgins DG, Heringa J. 2000. T-Coffee: A novel method for fast and accurate multiple sequence alignment. J Mol Biol. 302:205–217.

Pajares M, Jimenez-Moreno N, Dias IHK, Debelec B, Vucetic M, Fladmark KE, Basaga H, Ribaric S, Milisav I, Cuadrado A. 2015. Redox control of protein degradation. Redox Biol. 6:409–420.

Papetti C, Harms L, Windisch HS, Frickenhaus S, Sandersfeld T, Jurgens J, Koschnick N, Knust R, Portner HO, Lucassen M. 2015. A first insight into the spleen transcriptome of the notothenioid fish *Lepidonotothen nudifrons*: Resource description and functional overview. Mar Genomics 24:237–239.

Parker SK, Detrich HW. 1998. Evolution, organization, and expression of alpha-tubulin genes in the Antarctic fish Notothenia coriiceps - Adaptive expansion of a gene family by recent gene duplication, inversion, and divergence. J Biol Chem. 273:34358–34369.

Peck LS. 2016. A Cold Limit to Adaptation in the Sea. Trends Ecol Evol. 31:13–26.

Peck LS. 2018. Antarctic marine biodiversity: Adaptations, environments and responses to change. Oceanogr. Mar. Biol. Ann. Rev. 56: (in press)

Pe’er I, Felder CE, Man O, Silman I, Sussman JL, Beckmann JS. 2004. Proteomic signatures: Amino acid and oligopeptide compositions differentiate among phyla. Proteins Struct Funct Genet 54:20–40.

Place S, Hofmann G. 2005. Constitutive expression of a stress-inducible heat shock protein gene, hsp70, in phylogenetically distant Antarctic fish. Polar Biol. 28:261–267.

Place SP, Zippay ML, Hofmann GE. 2004. Constitutive roles for inducible genes: evidence for the alteration in expression of the inducible hsp70 gene in Antarctic notothenioid fishes. Am J Physiol Regul Integr Comp Physiol. 287:R429–R436.

Portner HO, Peck L, Somero G. 2007. Thermal limits and adaptation in marine Antarctic ectotherms: an integrative view. Philos Trans R Soc Lond B Biol Sci. 362:2233–2258.

Privalov PL. 1990. Cold denaturation of proteins. Biophys J. 57:A26–A26.

Pucci F, Rooman M. 2017. Physical and molecular bases of protein thermal stability and cold adaptation. Curr Opin Struct Biol. 42:117–128.

Pucciarelli S, Parker SK, Detrich HW, Melki R. 2006. Characterization of the cytoplasmic chaperonin containing TCP-1 from the Antarctic fish *Notothenia coriiceps*. Extremophiles 10:537–549.

Regoli F, Principato GB, Bertoli E, Nigro M, Orlando E. 1997. Biochemical characterization of the antioxidant system in the scallop *Adamussium colbecki,* a sentinel organism for monitoring the Antarctic environment. Polar Biol. 17:251–258.

Roach JC. 2002. A clade of trypsins found in cold-adapted fish. Protein Struct Funct Genet. 47:31–44.

Romero-Romero ML, Ingles-Prieto A, Ibarra-Molero B, Sanchez-Ruiz JM. 2011. Highly anomalous energetics of protein cold denaturation linked to folding-unfolding kinetics. PLoS One 6: e23050

Romisch K, Collie N, Soto N, Logue J, Lindsay M, Scheper W, Cheng CHC. 2003. Protein translocation across the endoplasmic reticulum membrane in cold-adapted organisms. J Cell Sci. 116:2875–2883.

Saunders NFW, Thomas T, Curmi PMG, Mattick JS, Kuczek E, Slade R, Davis J, Franzmann PD, Boone D, Rusterholtz K, et al. 2003. Mechanisms of thermal adaptation revealed from the genomes of the Antarctic Archaea *Methanogenium frigidum* and *Methanococcoides burtonii*. Genome Res. 13:1580–1588.

Scher HD, Whittaker JM, Williams SE, Latimer JC, Kordesch WEC, Delaney ML. 2015. Onset of Antarctic Circumpolar Current 30 million years ago as Tasmanian Gateway aligned with westerlies. Nature 523:580–583.

Shevenell AE, Ingalls AE, Domack EW, Kelly C. 2011. Holocene Southern Ocean surface temperature variability west of the Antarctic Peninsula. Nature 470:250–254.

Shin SC, Ahn DH, Kim SJ, Pyo CW, Lee H, Kim MK, Lee J, Lee JE, Detrich HW, Postlethwait JH, et al. 2014. The genome sequence of the Antarctic bullhead notothen reveals evolutionary adaptations to a cold environment. Genome Biol. 15:468.

Shin SC, Kim SJ, Lee JK, Ahn DH, Kim MG, Lee H, Lee J, Kim BK, Park H. 2012. Transcriptomics and Comparative Analysis of Three Antarctic Notothenioid Fishes. PLoS One 7: e43762.

Simão FA, Waterhouse RM, Ioannidis P, Kriventseva EV, Zdobnov EM. 2015. BUSCO: assessing genome assembly and annotation completeness with single-copy orthologs. Bioinform. 31:3210–3212.

Somero GN, Devries AL. 1967. Temperature tolerance of some Antarctic fishes. Science 156:257–258.

Stadtman ER, Moskovitz J, Berlett BS, Levine RL. 2002. Cyclic oxidation and reduction of protein methionine residues is an important antioxidant mechanism. Mol Cell Biochem. 234:3–9.

Struvay C, Feller G. 2012. Optimization to Low Temperature Activity in Psychrophilic Enzymes. Internat J Mol Sci. 13:11643–11665.

Szklarczyk D, Franceschini A, Wyder S, Forslund K, Heller D, Huerta-Cepas J, Simonovic M, Roth A, Santos A, Tsafou KP, et al. 2015. STRING v10: protein-protein interaction networks, integrated over the tree of life. Nucl Acids Res. 43:D447–D452.

Todgham AE, Crombie TA, Hofmann GE. 2017. The effect of temperature adaptation on the ubiquitin-proteasome pathway in notothenioid fishes. J Exp Biol. 220:369–378.

Todgham AE, Hoaglund EA, Hofmann GE. 2007. Is cold the new hot? Elevated ubiquitin-conjugated protein levels in tissues of Antarctic fish as evidence for cold-denaturation of proteins in vivo. Comp Biochem Physiol B Biochem System Environ Physiol. 177:857–866.

Uren Webster TM, Santos EM. 2015. Global transcriptomic profiling demonstrates induction of oxidative stress and of compensatory cellular stress responses in brown trout exposed to glyphosate and Roundup. BMC Genomics 16:32.

Vilella AJ, Severin J, Ureta-Vidal A, Heng L, Durbin R, Birney E. 2009. EnsemblCompara GeneTrees: Complete, duplication-aware phylogenetic trees in vertebrates. Genome Res. 19:327–335.

Windisch HS, Lucassen M, Frickenhaus S. 2012. Evolutionary force in confamiliar marine vertebrates of different temperature realms: adaptive trends in zoarcid fish transcriptomes. BMC Genomics 13:549.

Xu QH, Cheng CHC, Hu P, Ye H, Chen ZZ, Cao LX, Chen L, Shen Y, Chen LB. 2008. Adaptive evolution of hepcidin genes in Antarctic notothenioid fishes. Mol Biol Evol. 25:1099–1112.

Yang LL, Tang SK, Huang Y, Zhi XY. 2015. Low temperature adaptation is not the opposite process of high temperature adaptation in terms of changes in amino acid composition. Genome Biol Evol. 7:3426–3433.

Yang ZH. 2007. PAML 4: Phylogenetic analysis by maximum likelihood. Mol Biol Evol. 24:1586–1591.

Zerbino DR, Achuthan P, Akanni W, Amode MR, Barrell D, Bhai J, Billis K, Cummins C, Gall A, Giron CG, et al. 2018. Ensembl 2018. Nucl Acids Res. 46:D754–D761.

Zhao JS, Deng YH, Manno D, Hawari J. 2010. Shewanella spp. Genomic evolution for a cold marine lifestyle and *in-situ* explosive biodegradation. PLoS One 5:e9109

